# Carcinoma-associated fibroblast-like tumor cells remodel the Ewing sarcoma tumor microenvironment

**DOI:** 10.1101/2023.04.12.536619

**Authors:** Emma D. Wrenn, April A. Apfelbaum, Erin R. Rudzinski, Xuemei Deng, Wei Jiang, Sudha Sud, Raelene A. Van Noord, Erika A. Newman, Nicolas M. Garcia, Virginia J. Hoglund, Shruti S. Bhise, Sami B. Kanaan, Olivia G. Waltner, Scott N. Furlan, Elizabeth R. Lawlor

## Abstract

Tumor heterogeneity is a major driver of cancer progression. In epithelial-derived malignancies, carcinoma-associated fibroblasts (CAFs) contribute to tumor heterogeneity by depositing extracellular matrix (ECM) proteins that dynamically remodel the tumor microenvironment (TME). Ewing sarcomas (EwS) are histologically monomorphous, mesenchyme-derived tumors that are devoid of CAFs. Here we identify a previously uncharacterized subpopulation of transcriptionally distinct EwS tumor cells that deposit pro-tumorigenic ECM. Single cell analyses revealed that these CAF-like cells differ from bulk EwS cells by their upregulation of a matrisome-rich gene signature that is normally repressed by EWS::FLI1, the oncogenic fusion transcription factor that underlies EwS pathogenesis. Further, our studies showed that ECM-depositing tumor cells express the cell surface marker CD73, allowing for their isolation ex vivo and detection in situ. Spatial profiling of tumor xenografts and patient biopsies demonstrated that CD73^+^ EwS cells and tumor cell-derived ECM are prevalent along tumor borders and invasive fronts. Importantly, despite loss of EWS::FLI1-mediated gene repression, CD73^+^ EwS cells retain expression of EWS::FLI1 and the fusion-activated gene signature, as well as tumorigenic and proliferative capacities. Thus, EwS tumor cells can be reprogrammed to adopt CAF-like properties and these transcriptionally and phenotypically distinct cell subpopulations contribute to tumor heterogeneity by remodeling the TME.

## INTRODUCTION

Intratumoral heterogeneity promotes cancer growth, metastasis, therapeutic failure, and drug resistance (1, 2). Carcinoma-associated fibroblasts (CAFs) contribute to heterogeneity of epithelial-derived tumors by depositing extracellular matrix (ECM) proteins that dynamically remodel the tumor microenvironment (TME) (3). In addition, genomic instability of tumor cells leads to clonal evolution that contributes to mutational heterogeneity of the tumor cell compartment (2). While genomic instability can drive tumor cell heterogeneity, more recent studies have highlighted the role of non-mutational epigenetic heterogeneity (4). Single-cell sequencing and spatial analyses demonstrate that tumor ecosystems are comprised of heterogeneous tumor cells with distinct epigenetic and phenotypic states (5, 6). Non-mutational tumor cell heterogeneity is particularly relevant to pediatric tumors which typically have low mutation rates but high transcriptional and phenotypic heterogeneity due to hijacking of developmental and epigenetic regulators (7). Phenotypically distinct tumor cell subpopulations differentially contribute to tumor growth, metastatic potential, and therapy response, highlighting the critical importance of studying tumor cell subpopulations in the context of contextually relevant ecosystems (8–10).

Ewing sarcomas (EwS) are histologically primitive, monomorphous bone and soft tissue tumors that present across the lifespan, with a peak incidence in adolescence (11). Intensive cytotoxic therapy has improved prognosis for patients with localized tumors, but outcomes for patients with metastastic and relapsed EwS remain dismal (12). EwS tumors are initiated by chromosomal translocations between FET and ETS gene family loci, that generate chimeric in-frame fusion proteins, most commonly EWS::FLI1 (11). These fusions function as oncogenic transcription factors that cause widespread epigenomic reprogramming and aberrant activation and repression of hundreds of target genes (11, 13, 14). Recurrent mutations in other genes are uncommon and propagation of EwS tumors depends on continued expression of the fusion (11). Although intratumoral genomic instability is not a known feature of EwS, rare subpopulations of tumor cells that express low levels of EWS::FLI1 have been identified and these cells display enhanced metastatic properties in tumor models (15). In addition, modulation of EWS::FLI1-dependent gene signatures, without altering expression of the fusion itself, influences tumor cell state (reviewed in Ref. (16)). Genomic loss of *STAG2* (17, 18), expression of tissue-specific transcription factors (19–21), activation of the Wnt/β-catenin pathway (22), and other cell-intrinsic and cell-extrinsic mechanisms (16), all influence transcriptional activity of EWS::FLI1 and affect the tumorigenic and metastatic properties of EwS cells. Importantly, despite different underlying molecular origins, EWS::FLI1-depleted cells universally show increased expression of mesenchymal lineage genes that are normally repressed by the fusion. The contribution of EWS::FLI1-low, mesenchymal-high state tumor cells to EwS pathogenesis has yet to be defined in the context of tumors in vivo.

Here, we set out to identify and characterize EWS::FLI1-low cells in EwS tumors in situ. Transcriptomic profiling of EwS cell lines and patient tumors identified *NT5E* (CD73) as a cell surface marker that consistently marks EWS::FLI1-low tumor cells in vitro and in vivo. Phenotypic, proteogenomic, and immunofluorescence profiling confirmed that CD73^+^ cells exist as spatially distributed cell subpopulations that are enriched along tumor borders and invasive fronts. Further, our studies showed that CD73^+^ cells deposit protumorigenic ECM proteins such as tenascin-C (TNC), collagens, and proteoglycans, proteins that are largely deposited by CAFs in carcinomas (23). This matrisomal gene program is normally repressed by EWS::FLI1, validating that CD73^+^ tumor cells have lost EWS::FLI1 activity. Despite this, we found that expression of the EWS::FLI1-activated gene signature is largely retained by CAF-like tumor cells, revealing the novel discovery that the activating and repressive properties of the fusion can be dissociated in individual tumor cells, creating cells with hybrid transcriptional states. Together these studies identify the existence of transcriptionally and phenotypically distinct subpopulations of EwS tumor cells that contribute to tumor heterogeneity and remodel the TME via deposition of pro-tumorigenic ECM.

## RESULTS

### CD73 marks mesenchymal-high state EwS cell subpopulations

To study EWS::FLI1-low tumor cells in patient tumors and tumor xenografts in vivo, we first sought to define a cell surface marker that would permit their identification and isolation from bulk tumor cells. Given that EWS::FLI1 normally represses mesenchymal lineage genes (24, 25), we reasoned that expression of mesenchymal genes could be used to discriminate EWS::FLI1-low cells. Single cell transcriptomic studies recently showed that EwS cells exist along a transcriptional axis that partially reflects a mesenchyme development trajectory (19, 26). Using these data, we confirmed that activation of a mesenchymal development signature highly correlates with loss of EWS::FLI1-dependent gene repression (Figure 1A). Next, we intersected the two gene signatures to identify transcripts that are normally repressed by EWS::FLI1 (27) and expressed by mesenchymal-high state tumor cells (19) Fifty-two transcripts were included in the overlap, including *NT5E* which encodes the cell surface protein CD73 (Figure 1B). *NT5E*/CD73 (*5’-nucleotidase ecto)* is a GPI-anchored enzyme that is highly expressed by normal mesenchymal stem cells (MSC) (28), and we and others have previously shown that its expression is transcriptionally inhibited by EWS::FLI1 (19, 25).

**Figure 1.**
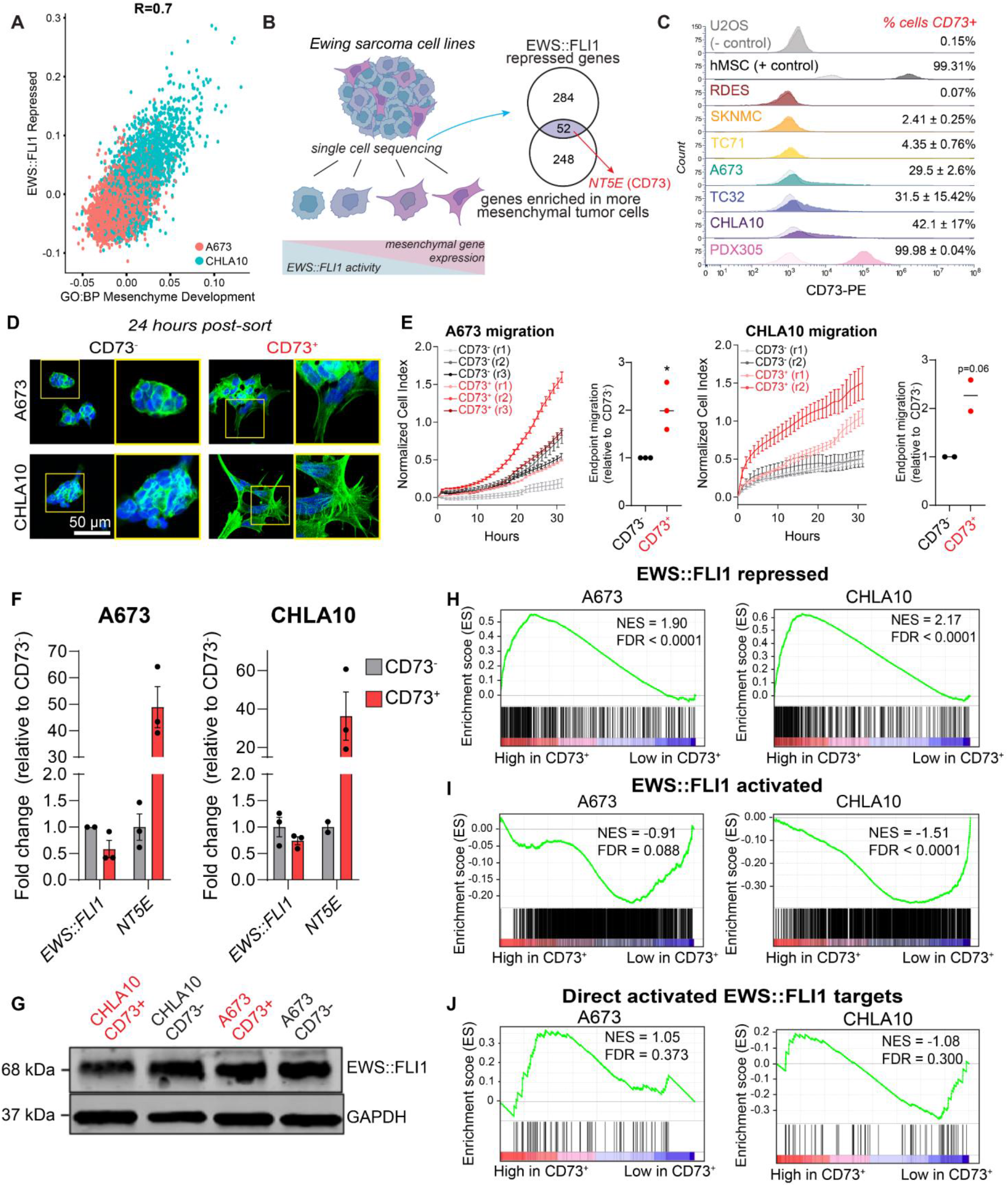
CD73 marks mesenchymal tumor cells that express EWS::FLI1-repressed genes. A) Scatter plot depicting expression of an EWS::FLI1 repressed geneset (27) and mesenchyme development genes (GO:0060485) in A673 and CHLA10 single-cell sequencing data (19). B) Left, Ewing sarcoma cells exist along a spectrum of mesenchymal identity. Genes associated with mesenchymal identity are repressed by EWS::FLI1. Right, Venn diagram depicting genes enriched in mesenchymal state EwS cells that are in both GO:BP Mesenchyme Development and EWS::FLI1-repressed (Kinsey et al. 2006) gene sets. C) Flow cytometry of CD73 cell surface expression in EwS and non-EwS (U2OS, hMSC) cell lines (CD73 = dark, isotype control = light). D) DAPI and F-actin staining of FACS-sorted CD73^-^ and CD73^+^ EwS cells in 2D culture, 24 hours after sorting (representative images of n=3). E) Left, real-time transwell cell migration assay of isogenic CD73^-^ and CD73^+^ cells (A673 n=3, CHLA10 n=2, r = biological replicate #, SEM of 3-8 technical replicates shown). Right, relative quantification of migration at endpoint for each replicate. F) RT-qPCR of *EWS::FLI1* and *NT5E* expression in CD73-sorted cell populations (n=2-3, error bars=SEM). G) Immunoblot of EWS::FLI1 protein expression in CD73-sorted cells. GAPDH is loading control. Representative of n=2. H-J) Gene set enrichment analysis (GSEA) of RNAseq data generated from CD73-sorted cell populations (n=3) H) genes repressed by EWS::FLI1 expression (27), I) genes activated by EWS:FLI1 expression (27), and J) 78 genes directly bound and upregulated by EWS::FLI1 (26).

Having identified CD73 as a potential marker of EWS::FLI1-low cells, we next performed flow cytometry on a panel of eight established and one early passage PDX-derived EwS cell lines. Human MSC and osteosarcoma cell line U2OS were used as positive and negative controls, respectively. All cells were grown in standard tissue culture conditions (see Methods section). Surface expression of CD73 by EwS cells was highly heterogeneous both within and between cell lines (Figure 1C). Cell lines ranged from fewer than 1% (RDES) to nearly 100% (PDX305) CD73^+^ cells. CD73^+^ fractions in the remaining EwS lines ranged from 2-42% and we observed remarkable reproducibility among biologic replicates demonstrating that each model grows in a preferred homeostatic and heterogeneous state. CD73^+^ cells were always present in large numbers in A673, CHLA10, and TC32 cultures (ranging from ∼15-50%), whereas CD73^+^ cells rarely comprised more than 5% of RDES, SK-N-MC, and TC71 (Figure 1C).

Next, we compared mesenchymal properties of CD73-high (>30% CD73^+^) and CD73-low (<5% CD73^+^) cell lines. As shown (Supplemental Figure S1A), CD73-low cell lines grew as rounded cells and formed multicellular clusters with few membrane protrusions. In contrast, cell lines with a high proportion of CD73+ cells were flattened and spindle-shaped with F-actin rich membrane protrusions (Supplemental Figure S1A). Cell lines with rare CD73^+^ cells were almost entirely non-invasive in 3D collagen I matrices, whereas CD73-high cell lines were robustly invasive (Supplemental Figure S1B). Fluorescence-activated cell sorting (FACS) was then used to isolate CD73^+^ and CD73^-^ cell fractions from the same A673 and CHLA10 cell cultures (Supplemental Figure S1C). Isolated CD73^-^ cell fractions grew as round cells that formed multicellular clusters with extensive cell-cell contacts and generated few membrane protrusions (Figure 1D). Conversely, syngeneic CD73^+^ cells displayed mesenchymal morphologies and generated extensive F-actin rich membrane protrusions (Figure 1D). In addition, sorted CD73^+^ cells migrated faster than corresponding CD73^-^ cells from the same culture (Figure 1E). Significantly, sorted cell populations reproducibly returned to their baseline mixed populations over one to two weeks, demonstrating that EwS cells are plastic and interconvert between CD73^+^ and CD73^-^ states (not shown). Thus, CD73 expression is remarkably plastic and heterogeneous both within and between EwS models and surface expression of CD73 reproducibly marks subpopulations of EwS cells that harbor mesenchymal properties.

### CD73^+^ EwS cells retain tumorigenicity and EWS::FLI1 expression

Having established that CD73 surface expression marks EwS cells that have lost EWS::FLI1-mediated repression of mesenchymal development genes, we next measured EWS::FLI1 to affirm that CD73^+^ cells are, in fact, EWS::FLI1-low cells. Unexpectedly, we detected no reproducible depletion of EWS::FLI1 transcript (Figure 1F) or protein (Figure 1G) in CD73^+^ vs. CD73^-^ cells. In addition, proliferation (Supplemental Figure S1D) and in vivo tumorigenic capacities (Supplemental Figure S1E) of FACS-sorted cells were equivalent or more pronounced in CD73^+^ than syngeneic CD73^-^ cells, corroborating the observation that CD73^+^ cells had not downregulated expression of the oncogenic fusion. To determine if the relative transcriptional activity of EWS::FLI1 was altered in CD73^+^ cells we performed RNA-seq on FACS-sorted cell populations of both A673 and CHLA10 models (Supplemental Table 1). This transcriptomic profiling confirmed that CD73^+^ EwS cells express high levels of the EWS::FLI1-repressed gene signature (Figure 1H). Surprisingly, however, this loss of EWS::FLI1-mediated gene repression was not accompanied by a corresponding loss of fusion-dependent gene activation (Figure 1I, J). CD73^+^ EwS cells displayed only modest loss of the EWS::FLI1-activation signature, no significant difference in expression of 78 direct EWS::FLI1 activated targets (26) (Figure 1I, J), and no loss of expression of the cell cycle program that is positively regulated by the fusion (Supplemental Figure S1F). Consistent with the observed mesenchymal phenotypes of CD73^+^ EwS cells, differential gene expression analysis identified the epithelial mesenchymal transition (EMT) signature as their most enriched hallmark signature compared to CD73^-^ cells (Supplemental Figure S1G). Thus, these studies together demonstrated that CD73 surface expression identifies subpopulations of tumor cells that have de-repressed the EWS::FLI1-repressed signature but that apparently retain expression of the fusion and much of its tumorigenic activation signature.

### CD73^+^ EwS tumor cells selectively upregulate expression of a pro-tumorigenic matrisomal gene program

The observation of potential dissociation between EWS::FLI1-activated and -repressed signatures in FACS-sorted populations of CD73^+^ EwS cells led us to more deeply investigate the relationship between phenotype and gene expression in individual cells. To achieve this, we exploited single cell proteogenomic profiling using CITE-seq (29). Transcriptomes and cell surface CD73 protein expression were measured in over 12,000 cells from eight established EwS cell lines and one early passage PDX-derived line (Figure 2A and Supplemental Figure S2A). Expression of *NT5E/*CD73 was profoundly heterogenous within and between models and a positive correlation between mRNA and surface protein expression suggested that the CD73 status of EwS cells is largely transcriptionally determined (Figure 2B, C). The transcriptomes of 1,760 *NT5E^+^* cells were next compared to 10,466 *NT5E^-^*cells across all cell lines and then in each cell line independently to identify markers of NT5E^+^ cells (Supplemental Table 2). As shown (Supplemental Figure S2B and Figure 2D), and consistent with data from RNA-seq of FACS-sorted cells, gene ontology analysis of differentially expressed genes identified the EMT gene signature as being reproducibly upregulated in *NT5E^+^*cells across all nine EwS models.

**Figure 2.**
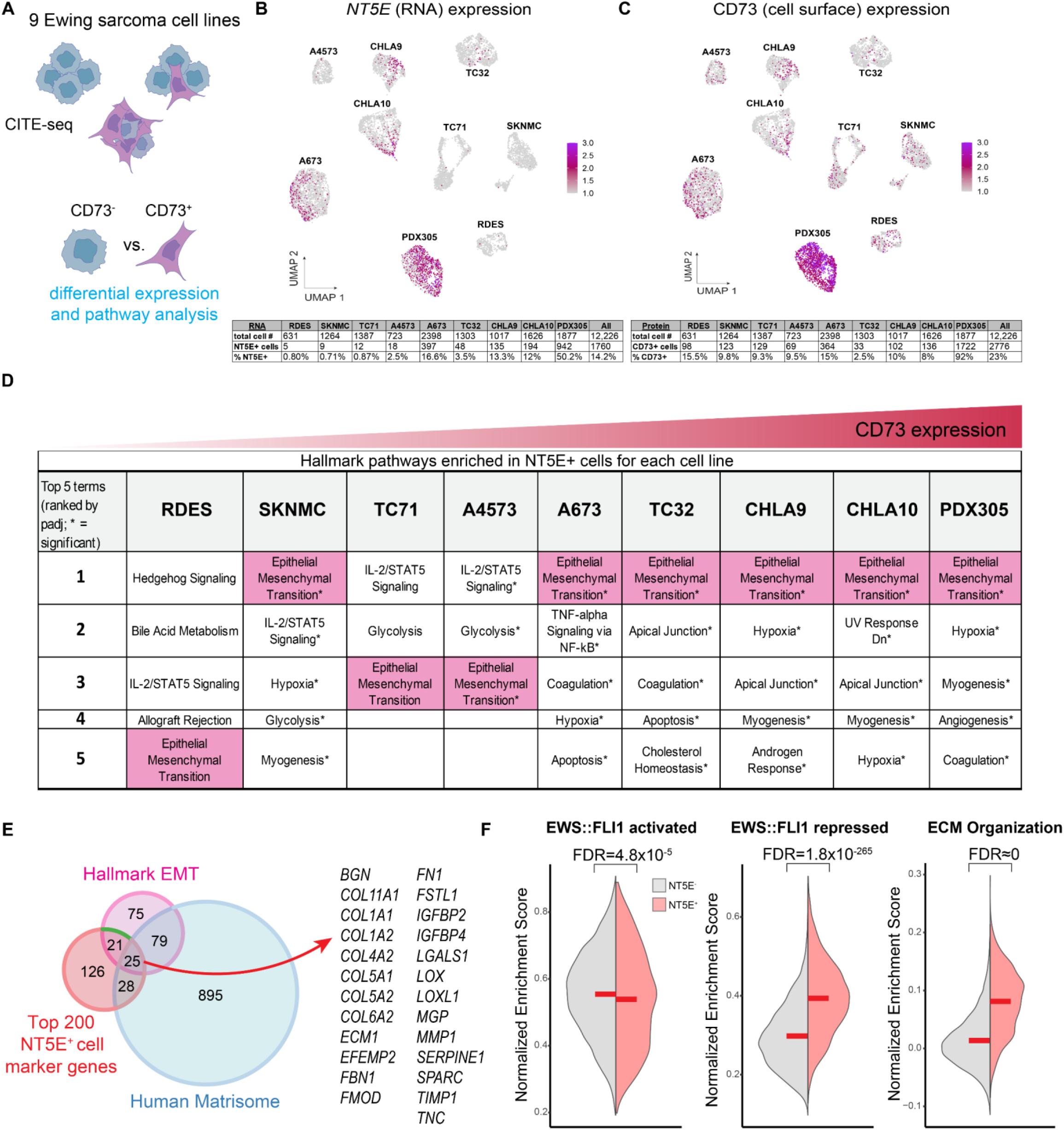
CD73+ EwS tumor cells selectively upregulate expression of a pro-tumorigenic matrisomal gene program. A) Workflow of CD73 CITE-seq and downstream differential expression analyses. B-C) UMAPs of CITE-seq data for *NT5E* (RNA) or CD73 (cell surface) expression in 9 EwS cell lines. Below, number of NT5E/CD73^+^ cells by cell line. D) Hallmark gene sets enriched in *NT5E^+^*cells from each cell line. E) Venn diagram overlap of Hallmark EMT genes, human matrisome genes (62), and the 200 top markers of *NT5E*+ cells as determined by single cell CITE-seq. Right, the 25 genes present in all 3 gene sets. F) Split violin plots depicting a normalized enrichment score for expression of EWS::FLI1 regulated (27) and GO:BP ECM Organization (GO:0030198) genesets in *NT5E^+^* and *NT5E^−^* cells (CITE-seq data inclusive of 9 EwS cell lines).

EMT is a complex and highly conserved developmental process that is frequently hijacked during carcinoma pathogenesis (30). To better understand how an EMT signature could be differentially activated in sarcoma cells that are, by definition, mesenchymal (31), we explored the identity of the EMT signature genes that were upregulated in CD73^+^ cells. Forty-six of 200 EMT genes were differentially upregulated in *NT5E^+^*vs. *NT5E^-^* cells and we noted that many of them encode for ECM and matrisome-associated proteins (Figure 2E, Supplemental Figure S2C). These included numerous collagens, integrins, fibrillin 1 (*FBN1*), fibronectin (*FN*), secreted protein acidic and cysteine rich (*SPARC*), and tenascin-C (*TNC*) (Supplemental Figure S2D). Quantitative RT-PCR of FACS-sorted cells validated higher expression of ECM-related genes in CD73^+^ cell populations (Supplemental Figure S2E), and immunofluorescence confirmed increased deposition of TNC protein (Supplemental Figure S2F). Thus, these findings suggested that the transcriptional and phenotypic properties of CD73^+^ EwS tumor cells are similar to ECM-producing CAFs (3, 32).

Having established that the mesenchymal-high state of CD73^+^ EwS cells is characterized by significant and reproducible upregulation of matrisomal gene programs, we next evaluated expression of EWS::FLI1-activated and -repressed gene signatures in these cells. Consistent with our observations in CD73^+^ vs. CD73^-^ sorted cell populations (Figure 1H-J), single *NT5E*^+^ cells showed only a modest decrease in expression of the EWS::FLI1-activated signature, but a marked upregulation of EWS::FLI1-repressed and ECM genes (Figure 2F). These findings demonstrate a partial decoupling of EWS::FLI1-dependent gene activation and gene repression in *NT5E^+^* cells, with a significant and reproducible re-activation of mesenchymal and ECM genes that are normally repressed by the fusion.

### Spatial and transcriptional heterogeneity of EwS tumor cells exists in vivo

Given our finding of extensive intratumoral heterogeneity in vitro, we next investigated transcriptional and phenotypic profiles of EwS tumor cell states in vivo. Local and metastatic CHLA10 tumor xenografts were generated in immunocompromised mice by subcutaneous and tail vein injections, respectively. Tumors were harvested at humane endpoints and subjected to H&E staining and immunofluorescence (Figure 3A-F). As shown, CD73^+^ tumor cells were detected in all tumors with a notable increase in frequency in metastatic tail vein-derived tumors compared to subcutaneous masses (Figure 3B-F). Spatial heterogeneity of CD73^+^ cell subpopulations was also apparent within each tumor (Figure 3). In the tail vein-derived tumors, CD73^+^ cells were enriched in peri-necrotic regions, tumor borders, and in invasive foci that had infiltrated the retroperitoneal cavity (Figure 3E, F). In keeping with our in vitro findings, tumor cell-derived ECM deposition was evident in regions adjacent to the CD73^+^ tumor cells (Figure 3B-F). Similar spatial heterogeneity in CD73 expression and proximity to ECM deposition was observed in additional A673 and PDX305 subcutaneous xenografts (Supplemental Figure S3A, B).

**Figure 3.**
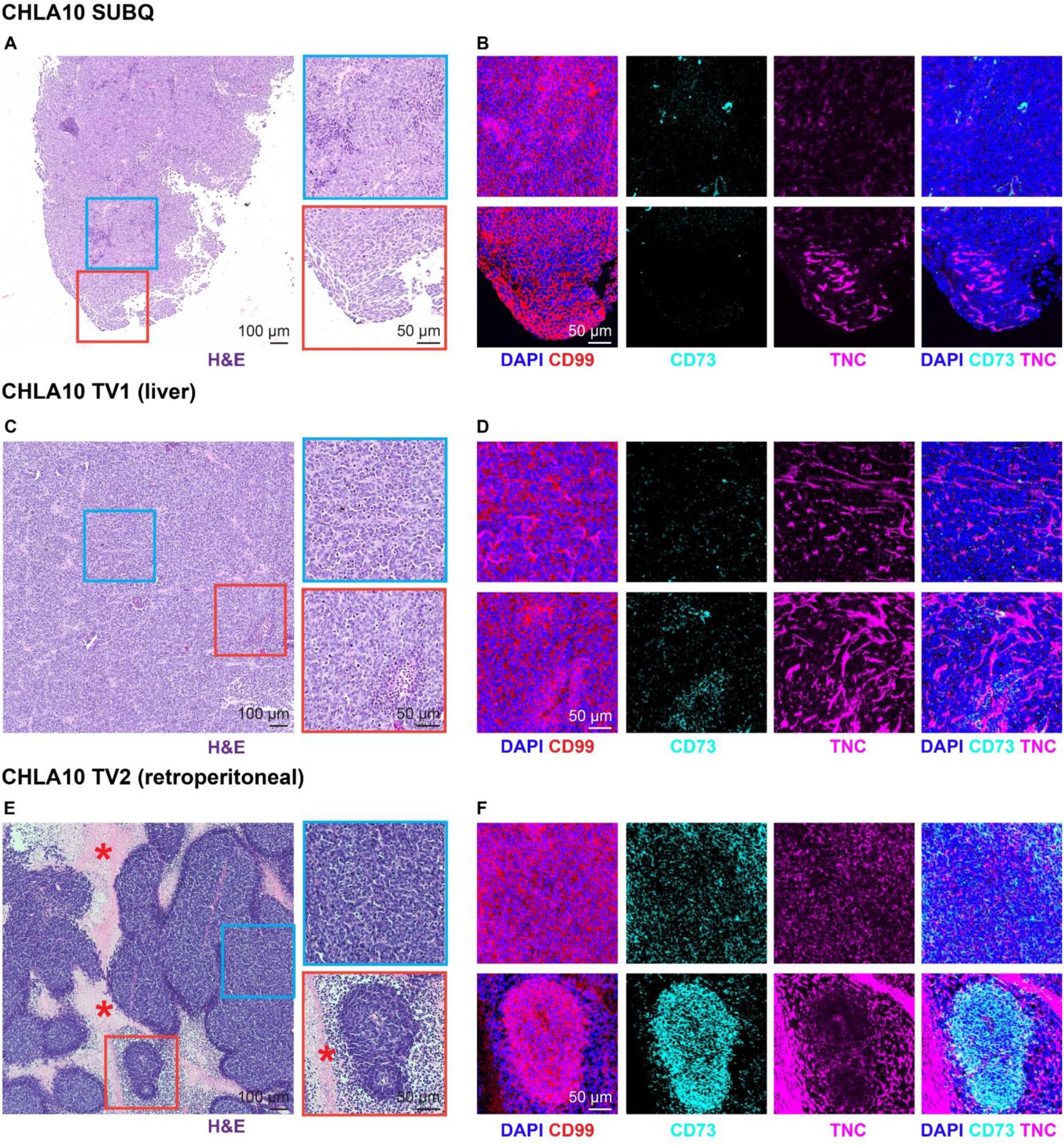
Spatial heterogeneity of CD73^+^ tumor cells and ECM deposition in vivo. FFPE sections were generated from 3 CHLA10 tumor xenografts from 3 separate mice: one subcutaneous tumor (SUBQ), a liver tumor (TV1), and a retroperitoneal tumor (TV2) formed after tail vein injection. A) H&E staining of the SUBQ xenograft tumor. Right, zoomed insets. B) Immunofluorescence of the same regions in adjacent SUBQ FFPE sections for DAPI (nuclei), CD99 (tumor membrane marker), CD73, and human TNC. C) H&E staining of the TV1 xenograft liver tumor. Right, zoomed insets. D) Immunofluorescence of the same regions in adjacent TV1 FFPE sections for DAPI (nuclei), CD99 (tumor membrane marker), CD73, and human TNC. E) H&E staining of the TV2 xenograft retroperitoneal tumor. Right, zoomed insets. Asterisk marks regional necrosis adjacent to viable tumor borders. F) Immunofluorescence of the same regions in adjacent TV2 FFPE sections for DAPI (nuclei), CD99 (tumor membrane marker), CD73, and human TNC.

We next performed integrated digital spatial profiling and whole human transcriptome analysis to define the transcriptional states of phenotypically distinct tumor cells in these in vivo tumor microenvironments (Figure 4A). Ki67 and cytoskeletal protein vimentin were used as morphology markers to identify differentially proliferative and mesenchymal regions, respectively(Figure 4B). We then selected four spatially distinct regions of interest (ROIs) from each tumor for capture and next generation sequencing (NGS). Between 1400 and 6500 cells were captured from each of the 12 ROIs which were distributed in tumor interiors (e.g. ROI1, ROI5, ROI11), along tumor-stromal borders (e.g. ROI2, ROI7), and in invasive tumor foci that were surrounded by necrotic tissue (e.g. ROI9, ROI10) (Figure 4B). Normalized *MKI67* and *VIM* gene expression in the captured ROIs (Supplemental Table 3) corroborated the observed spatial differences in Ki67 and vimentin immunofluorescence (Supplemental Figure S4A, B). As shown, tumor cells that were captured from subcutaneous and tail vein-derived focal liver tumors generally expressed higher levels of *MKI67* than cells captured from an infiltrating retroperitoneal tumor (Supplemental Figure S4A, B). In contrast, expression of *VIM* was generally higher in tail vein-derived metastatic tumors and heterogeneity among ROIs was apparent in all three tumors (Supplemental Figure S4A, B).

**Figure 4.**
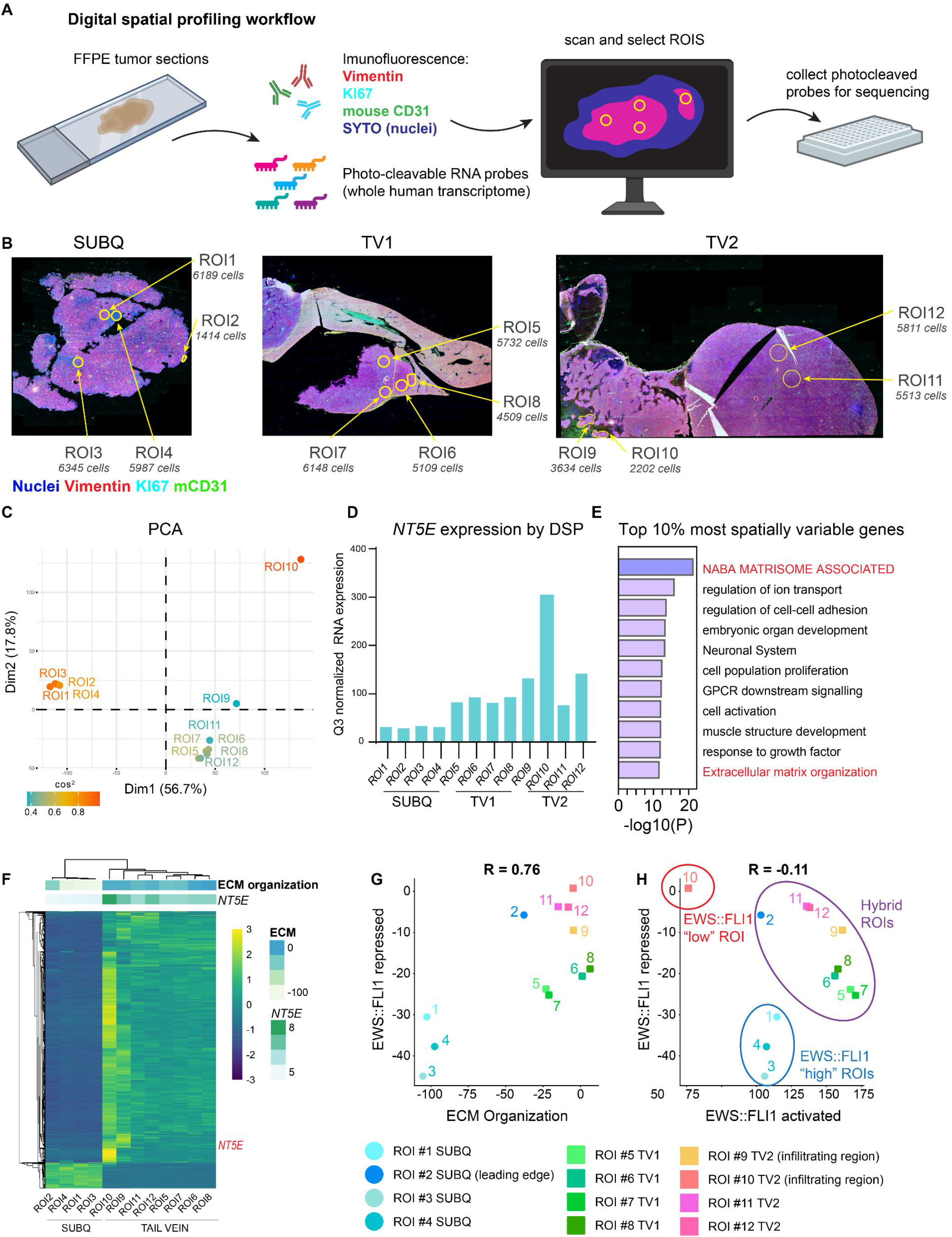
Digital spatial profiling reveals transcript and protein-level spatial heterogeneity in EwS tumors. A) Workflow of the Digital spatial profiling (DSP) method used. Briefly, immunofluorescence of Vimentin (mesenchymal cytoskeletal marker), KI67 (proliferative nuclei), SYTO (all nuclei), anti-mouse CD31 (murine blood vessels) were used to select regions of interest (ROIs) on xenograft sections hybridized with a whole human transcriptome probe library. ROIs were captured and gene expression quantified by next-generation sequencing after UV cleavage of probes from selected ROIs. B) Vimentin, KI67, mCD31, SYTO (nuclei) immunofluorescence and selected ROIs indicated for each of 3 tumors subjected to DSP: CHLA10 (SUBQ) and tail vein-derived liver (TV1) and retroperitoneal (TV2) tumors as in Figure 3. Approximate DAPI+ cell count is listed for each ROI. C) PCA plot of normalized whole human transcriptome gene expression for each ROI. Colored by cos^2^ value (indicates quality of representation by the principal components). D) Bar graph depicting Q3 normalized mRNA expression of *NT5E* within each ROI as determined by DSP and next generation sequencing. E) Gene ontology (Metascape) of the top 10% most spatially variable genes across 12 ROIs (highest coefficient of variation between ROIs, n=1339 genes). E) Unsupervised heatmap of the top 10% most spatially variable genes across all ROIs. Top panels show expression of *NT5E (*log_2_) and the GO:BP ECM Organization gene set. G) Scatter plot of expression of EWS::FLI1-repressed gene signature (27) vs. ECM Organization gene set expression. Individual ROIs are labeled. H) Scatter plot of RNA expression of EWS::FLI1 repressed signature vs. EWS::FLI1 activated signature in each ROI (27). EWS::FLI1-high, EWS::FLI1-low, and EWS::FLI1-hybrid ROIs are indicated.

Having confirmed that CHLA10 cells display heterogeneity of *VIM* expression in vivo, we next performed unsupervised analyses of whole transcriptomic data to interrogate spatial and biologic heterogeneity in an unbiased fashion. First, principal components analysis (PCA) of the 12 ROI-transcriptome dataset showed that subcutaneous tumor ROIs clustered tightly and separately from ROIs captured from metastatic tumors despite the fact that all tumors were generated from CHLA10 cells (Figure 4C). Transcriptomes of ROIs captured from tail vein-derived tumors also demonstrated considerably more heterogeneity than subcutaneous ROIs. This was especially notable when comparing tumor cells captured from tumor interiors to those captured from invasive tumor foci (ROI9 and ROI10) (Figure 4C). The transcriptomic profile of tumor cells in ROI10 was distinct from all other ROIs and, significantly, targeted analysis of the data reveals that these tumor cells expressed the highest level of *NT5E* (Figure 4D). Next, we sought to identify genes with the most spatial heterogeneity. To achieve this, we again took an unbiased approach and identified the most variably expressed genes across all 12 ROIs as determined by coefficient of variance (33). Limiting analysis to the top 10% most differentially expressed transcripts generated a list 1,339 genes and gene ontology analysis revealed these to be highly enriched for matrisomal and ECM-related gene programs (Figure 4E). Unsupervised hierarchical clustering of these 1,339 genes clearly separated subcutaneous from metastatic ROIs and showed that *NT5E* and an ECM organization gene signature are markedly upregulated in tumor cells captured from metastatic foci (Figure 4F; ROI9 & ROI10). One notable exception to this trend was ROI2, a region along the tumor: stromal interface of the subcutaneous tumor that highly expressed ECM genes but not *NT5E/*CD73. Thus, intratumoral spatial heterogeneity of EwS cell phenotypes and transcriptional profiles is evident in vivo. *NT5E* and matrisomal gene programs are, in general, more highly expressed by tumor cells in metastatic sites than those that are growing in localized masses and expression of ECM-encoding genes is particularly enriched along tumor borders and invasive foci.

Finally, we compared expression of EWS::FLI1-activated and -repressed genes in tumor cells captured from the 12 spatially distributed ROIs. As predicted from our in vitro data, EwS cells that highly expressed the ECM gene signature showed loss of EWS::FLI1-dependent gene repression (Figure 4G). However, loss of the repressive signature was not always accompanied by loss of EWS::FLI1-mediated gene activation, and hybrid transcriptional states were uncovered (Figure 4H). Transcriptomes of EwS cells isolated from the core of the subcutaneous tumor (ROIs 1, 3, & 4) were typical of EWS::FLI1-high cells (activated genes ON/repressed genes OFF). Likewise, the ROI10 transcriptional signature was fully compatible with EWS::FLI1-low cells (activated genes OFF/repressed genes ON). In contrast, transcriptomes of tumor cells captured from the remaining regions displayed features of both EWS::FLI1-high and -low cells. This is particularly evident for ROIs 9,11, and 12 wherein EwS cells were found to express high levels of both EWS::FLI1-activated and -repressed genes. Thus, these studies demonstrate dissociation of EWS::FLI1 activating and repressive transcriptional activity in EwS tumor regions in vivo and suggest that tumor subpopulations can upregulate ECM signatures without downregulating expression of EWS::FLI1-activated genes. Although it is possible that hybrid ROIs were comprised of mixed populations of EWS::FLI1-high and EWS::FLI1-low cells, when combined with our identification of transcriptionally hybrid states in vitro (Fig 2), these data lead us to conclude that hybrid EwS cells are present in tumors in vivo in and these cells acquire mesenchymal properties while retaining EWS::FLI1-dependent gene activation.

### Heterogeneity of *NT5E/*CD73+ tumor cells and tumor-derived ECM is evident in patient tumors

There is growing evidence that cells in transitional EMT cell states, rather than fully epithelial or mesenchymal cells, drive growth and metastatic progression of carcinomas (34, 35). Having identified *NT5E/*CD73 as a marker of transcriptionally hybrid and phenotypically distinct CAF-like tumor cell subpopulations in EwS models, we sought to investigate the presence of these cells in primary patient tumors. Analysis of three EwS gene expression datasets (GSE34620, GSE142162, GSE17679) (18, 36, 37) revealed marked inter-tumor variability in *NT5E* expression (Figure 5A). Next, we identified genes that were strongly correlated with *NT5E* (R>0.6) across patient tumors and discovered a striking overlap among the independent cohorts (Figure 5B). Hallmark EMT and ECM-related gene ontology signatures were highly enriched in these 217 shared *NT5E*-correlated genes (Supplemental Table 4), corroborating the enrichment of these signatures in *NT5E*+ EwS cells in tumor models (Figure 5C). Integrating the top 200 markers of *NT5E+* EwS cells in vitro (Figure 2E, Supplemental Table 2) with the 217 genes *NT5E*-correlated genes in patient tumors in vivo, generated a list of 27 genes that were highly reproducibly upregulated alongside *NT5E* in EwS tumor cells and these include numerous EMT and ECM genes, most of which are normally repressed by EWS::FLI1 (27, 38, 39) (Figure 5D). Consistent with this observation, expression of the 28 gene signature by individual patient tumors correlated positively with expression of both ECM organization and EWS::FLI1-repressed gene signatures (Figure 5E and Supplemental Figure S5A-B).

**Figure 5.**
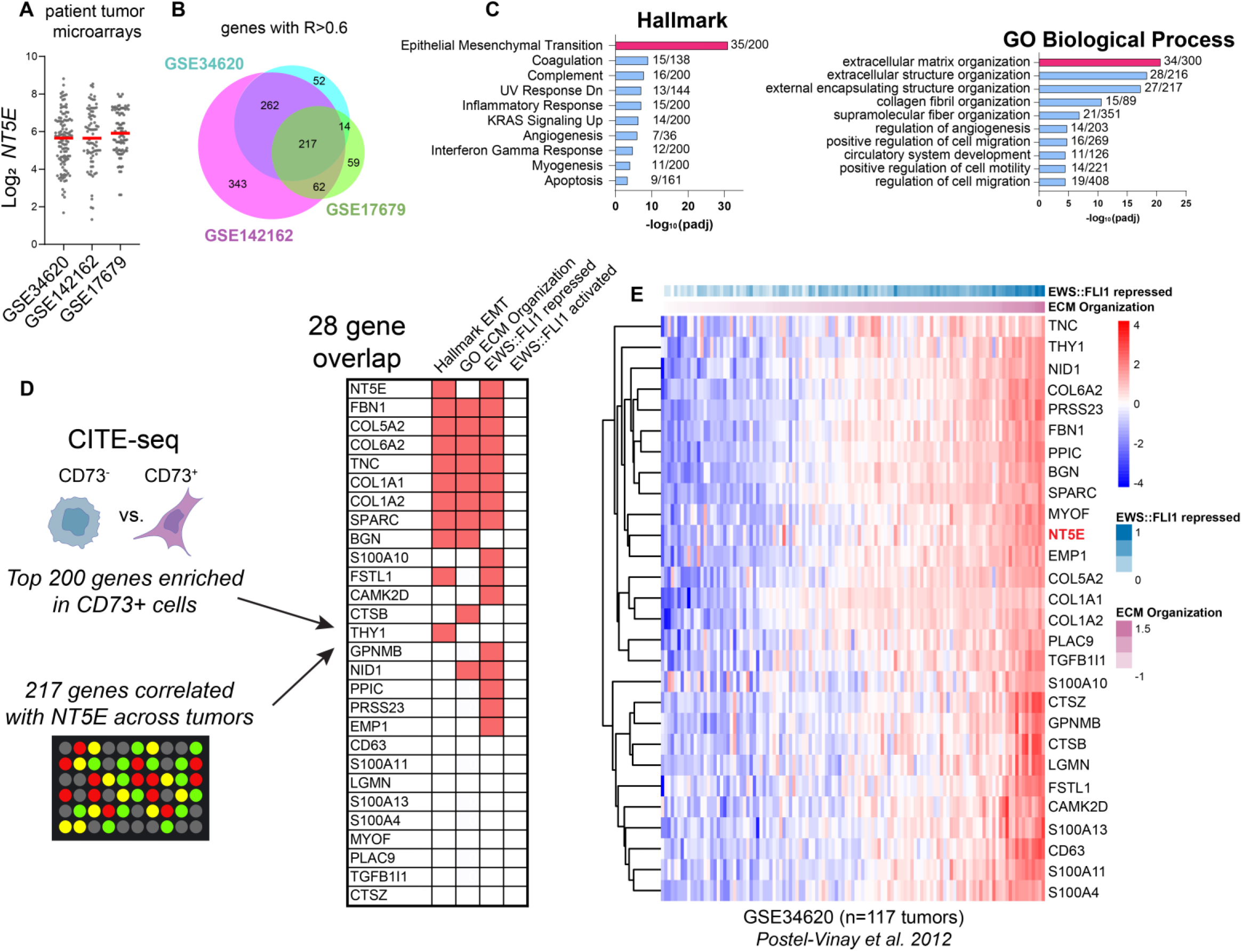
Heterogeneity of *NT5E* and related mesenchymal transcriptomic signatures in EwS patient tumor biopsies. A) Log_2_ *NT5E* expression derived from three independent gene expression microarray studies of patient tumors (18, 36, 37). B) Venn diagram of genes positively correlated with *NT5E* (R>0.6) in each dataset and the intersection of these genes. C) Hallmark and GO:BP gene ontology (Enrichr) of the 217 *NT5E*-correlated genes (and *NT5E* itself) shared across three studies. D) Workflow and characterization of the overlap between genes marking *NT5E*^+^ cells (from Figure 2) and genes highly correlated with *NT5E* in patient tumor biopsies. Right, membership of overlapping 28 genes in Hallmark EMT, GO:BP ECM Organization, EWS::FLI1 repressed (27, 38, 39), and EWS::FLI1 activated genesets (26, 27, 38, 39). E) Heatmap of *NT5E* and 27 associated marker gene expression vs. EWS::FLI1 repressed genes (27), ranked by ECM organization expression in Ewing sarcoma patient tumor microarray GSE34630 (36).

This analysis of bulk tumor RNA profiling confirmed inter-tumor heterogeneity of *NT5E* expression and the relationship between high *NT5E* and expression of EWS::FLI1-represssed matrisomal genes. To assess intratumoral heterogeneity and the potential existence of distinct CAF-like tumor cell subpopulations we evaluated CD73 and ECM protein expression in tumor sections from archived patient biopsies. We first stained a tumor tissue microarray (TMA) with an anti-CD73 antibody using the EWS::FLI1-induced protein NKX2-2 as a tumor marker. CD73^+^ cells were detected in only 7 of 24 tumor cores (20 unique patients; Supplemental Table 5) and the frequency of CD73+ tumor cells varied from rare to extensive (Supplemental Figure S6A-C). Staining of adjacent sections of the TMA with an antibody against TNC confirmed inter-and intra-tumoral heterogeneity of TNC and deposition of this protumorigenic ECM protein was most robust in regions adjacent to CD73^+^ tumor cells (Supplemental Figure S6D).

Given the observed spatial heterogeneity of CD73^+^ tumor cells and tumor-derived ECM in EwS xenografts (Figure 3, Supplemental Figure S3) and TMA cores (Supplemental Figure S6B, C), we next stained full biopsy sections from two tumors that were included in the TMA. One of these (#20) showed extensive CD73^+^ cells on the TMA core while the other (#35) was scored negative (Supplemental Table 5). Whole sections of both biopsies showed the presence of large acellular regions that denoted areas where core punches had been previously harvested for inclusion on TMAs (Figure 6A,B). Staining for the CD99 tumor marker showed that biopsy #20 included EwS tumor embedded within adjacent non-tumor tissue (Figure 6A) while biopsy #35 was predominantly comprised of viable tumor (Figure 6B). Importantly, CD73^+^ tumor cells within CD99^+^ regions were detected in both tumors, including biopsy #35 which was devoid of CD73^+^ cells on the TMA (Figure 6A,B). Spatial heterogeneity of ECM proteins TNC and biglycan (BGN) was also evident between and within the tumors and both proteins were more highly expressed adjacent to CD73^+^ tumor cells (Figure 6A,B). Thus, EwS tumor cells are both transcriptionally and phenotypically heterogeneous in vitro and in vivo and spatial co-localization of CD73^+^ tumor cells with pro-tumorigenic ECM suggests that EWS::FLI1-low and EWS::FLI1-high/low hybrid tumor cells function as CAF-like cells (Figure 6C).

**Figure 6.**
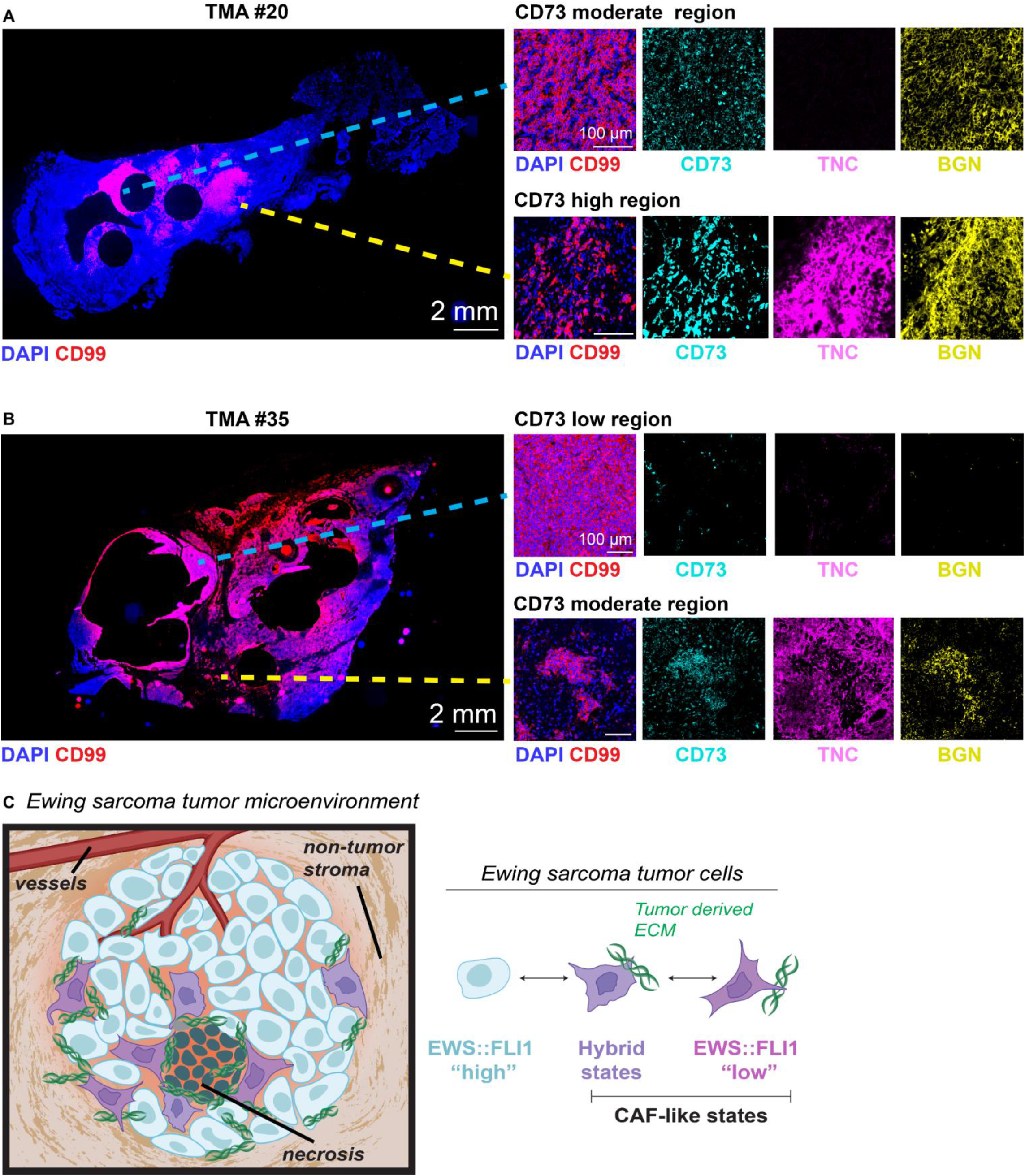
Intra-tumoral heterogeneity of CD73^+^ tumor cells and ECM deposition in patient biopsies. Immunofluorescence analysis of full tumor biopsy sections: (A) TMA ID #20 (extensive CD73+ scoring on TMA) and (B) TMA ID #35 (negative CD73 scoring on TMA). DAPI (nuclei), CD99 (tumor membrane marker), CD73, Tenascin-C, and biglycan. Right panels, zoomed insets show regions from the same tumor section with disparate CD73 expression and ECM deposition. C) Model of cell state heterogeneity in EwS wherein tumors contains spatially and transcriptionally heterogeneous subpopulations of tumor cells. Selective derepression of EWS::FLI1-suppressed mesenchymal target genes leads to acquisition of mesenchymal identity without loss of EWS::FLI1-dependent gene activation and tumorigenicity. These transcriptionally hybrid cells display properties of CAFs and promote remodeling of the TME by depositing pro-tumorigenic ECM.

## DISCUSSION

Accruing evidence suggests that tumor cell heterogeneity is a universal feature of all cancers and that even genetically homogeneous tumors develop extensive cell diversity through epigenetic processes (4, 5). Phenotypically distinct tumor cell subpopulations can contribute in different ways to tumor progression as they may have very different proliferation rates, metastatic potential, and differential responses to treatment (5). EwS are genetically quiet tumors that are driven by a single fusion oncoprotein (11). They show little evidence of clonal genetic diversity or evolution and they are histologically classified as homogeneous, small round blue cell tumors (12). Here, we have identified that EwS tumor cells exist in a previously uncharacterized spectrum of cell states that differentially express mesenchymal gene programs. In particular, our findings show that transcriptionally distinct subpopulations of highly mesenchymal tumor cells are present in EwS cell lines in vitro, xenograft models in vivo, and in patient tumors in situ. These cells can be identified by cell surface expression of CD73 and they are distinguished by their upregulation of matrisomal protein-encoding genes and their deposition of ECM. The distinct capacity of these cells to remodel the local TME through deposition of pro-tumorigenic ECM leads us to label them “CAF-like” EwS cells (Figure 6C).

The critical importance of ECM remodeling to tumor growth, invasion, and metastatic progression is well established (40–42). In epithelial-derived tumors ECM is largely deposited by CAFs, non-malignant stromal cells that most often arise from resident tissue fibroblasts or other mesenchymal cells and that are activated to secrete pro-tumorigenic matrisomal proteins in response to cues from the tumor cell compartment (3). More rarely, subpopulations of carcinoma cells themselves acquire the capacity to deposit ECM and promote pro-tumorigenic remodeling of the local TME and there is evidence that tumor cell-derived ECM confers a more aggressive tumor state (42, 43). The critical contribution of ECM and non-malignant stromal cells to sarcomas is less well studied but post-translational remodeling of collagen by tumor cells plays an essential role in mediating metastatic progression of soft tissue sarcomas (44). The contribution of ECM remodeling to growth and progression of EwS has not been deeply investigated, though clinico-pathologic correlative studies suggested that a stroma-rich TME influences EwS gene expression and patient prognosis (45). EwS cells and tumors can deposit fibrillar and non-fibrillar proteins (46–48) and we previously showed that activation of canonical Wnt/beta-catenin and TGF-beta signaling upregulates secretion of pro-tumorigenic and – angiogenic matrisomal proteins by EwS cells, including collagens and TNC (49, 50). In the current work, we have identified discrete subpopulations of EwS cells that deposit pro-tumorigenic ECM proteins in vivo. These cells are largely CD73^+^ and they are enriched on tumor borders, perinecrotic regions, and in invasive foci. Importantly, these cells were rarely found on our examination of small TMA tumor cores. Diagnostic EwS tumor specimens are normally obtained from needle or small open biopsies of viable tumor interiors (51). Thus, these CD73^+^ tumor cells and tumor cell-derived ECM can be missed when histologic and molecular analyses are limited to small tumor specimens. As our understanding of the critical importance of tumor heterogeneity and tumor: TME crosstalk grows, so too does our appreciation that TMA cores may not be adequate for studies of tumor biology that aim to dissect tumors as spatially dynamic ecosystems (52).

Our single cell studies of EwS cell transcriptomes and spatial profiling studies of tumors in vivo generated the unanticipated discovery that, although they expressed high levels of matrisomal genes that are normally repressed by EWS::FLI1, CD73^+^ EwS cells retained expression of the fusion and were largely not “EWS::FLI1-low”. While we did identify invasive cell clusters that were consistent with a fully EWS::FLI1-low state (e.g., Figure 4H, ROI10), CAF-like EwS cells most often existed in hybrid EWS::FLI1 transcriptional states. These cells retained expression of the fusion-activated gene signature alongside high expression of genes that are normally repressed by EWS::FLI1. Thus, these EwS cells gained mesenchymal properties, including the capacity to deposit ECM, without losing proliferative or tumorigenic potential. Notably, we also observed transcriptional heterogeneity among CAF-like populations in vivo, recapitulating the heterogeneity of CAFs that has been described in carcinomas (53). These observations lead us to conclude that EwS cell states are more nuanced than first appreciated. While more migratory and metastatic EWS::FLI1-low cells were first presumed to express lower levels of the fusion (15), it is now evident that the activity of the fusion can be influenced by cell intrinsic and extrinsic factors that modulate its transcriptional output (16). Our current work adds further complexity, revealing that the activating and repressing properties of the fusion can be decoupled in individual tumor cells. The molecular mechanisms that mediate this dissociation remain to be elucidated and this is a key gap in knowledge that the field must now fill. However, given the exciting work from several labs that cell context-specific transcription factors (19–21), chromatin remodelers (17, 18), and nuclear condensates (14, 54) influence EWS:FLI1 activity, it is likely that epigenetic mechanisms will be identified as drivers of these hybrid transcriptional states.

Tumor cell heterogeneity and cell plasticity are key factors in tumor evolution (4) and metastatic progression of carcinoma models is, at least in part, driven by transitional or hybrid cells that exist along a continuum between epithelial and mesenchymal states (34, 35). In addition, interactions between dynamic tumor cells and an equally dynamic TME create complex tumor ecosystems that support tumor growth, enable metastatic progression, and promote resistance to current treatment approaches (55–57). Given this, the tumor: TME interface and ECM remodeling processes are increasingly being investigated as novel targets on which to focus basic and translational cancer research efforts (23, 58–61). Here we have reported the existence of CAF-like EwS cells that contribute to ECM remodeling, and we propose that these cells are likely to play critical roles in mediating tumor progression and influencing treatment response. A deeper understanding of CAF-like EwS cells is now needed, including determination of their prevalence, origins, fundamental biology, and impact on neighboring tumor cells as well as the immune and non-immune TME. Elucidating the dynamic and highly heterogeneous nature of EwS ecosystems will afford novel insights into tumor biology that will guide development and rational integration of TME-targeted therapeutics into EwS treatment regimens.

## FIGURE LEGENDS

**Supplemental Figure S1.**
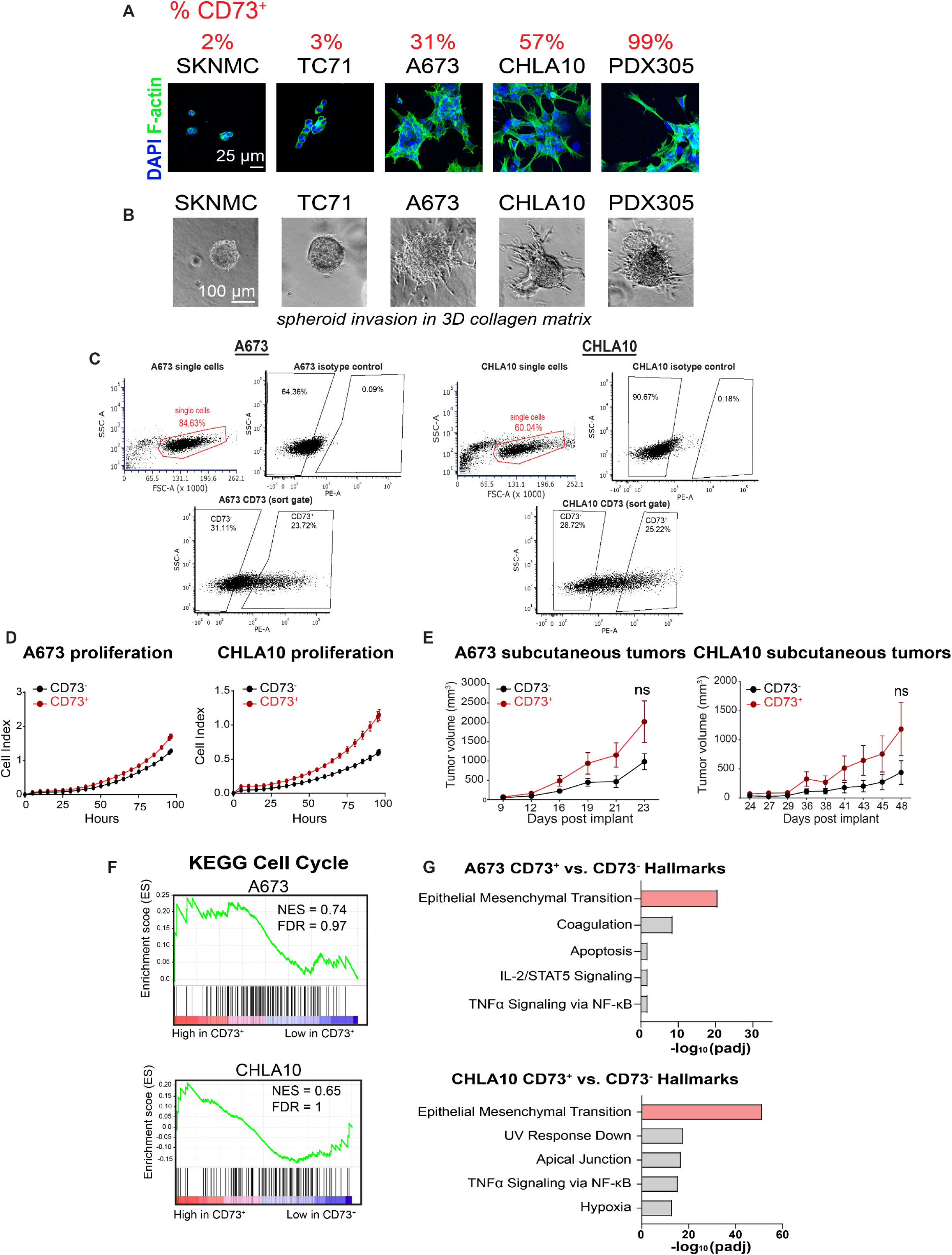
CD73+ EwS cells retain proliferative and tumorigenic capacity. Related to Figure 1. A) DAPI and F-actin staining of EwS cell lines in 2D culture. Panels ordered left to right based on CD73^+^ cell frequency. B) EwS cell spheroids from the same cell lines after 4 days in 3D spheroid culture in rat tail collagen. C) Flow cytometry gating strategy for CD73 FACS. D) Real-time proliferation assay of CD73^-^ vs. CD73^+^ sorted cells (n=1). Error bars = SEM of technical replicates. E) Subcutaneous tumor growth of CD73^-^ and CD73^+^ sorted cells implanted in NSG mice (n=5). *p*-values = unpaired t-tests. Error bars = SEM. F) GSEA on RNA-seq data from CD73^+^ vs. CD73^-^ cells for cell-cycle related genes (KEGG) (n=3). G) Gene ontology (Enrichr) of RNA-seq data for Hallmark gene sets in CD73^+^ vs. CD73^-^ cells.

**Supplemental Figure S2.**
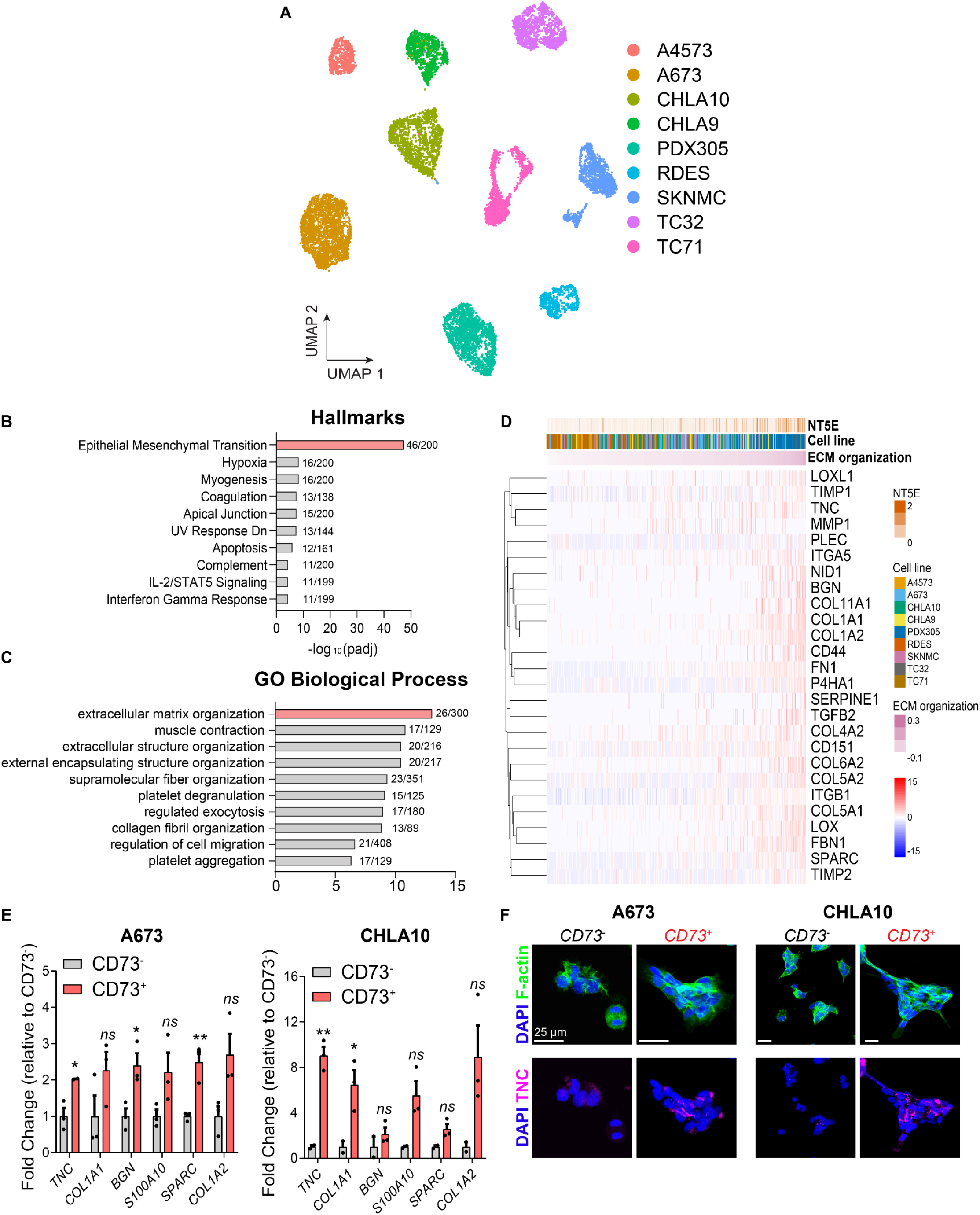
CD73+ cells are enriched in EMT and ECM gene expression. Related to Figure 2. A) UMAP of CITE-seq data showing 9 EwS cell lines. B-C) Gene ontology (Enrichr) of Hallmark gene sets (B) and GO Biological Process (C) of the top 200 significantly differentially expressed genes in NT5E^+^ vs. NT5E-cells (pooling 9 cell lines). D) Heatmap of expression (by single cell sequencing) of 26 ECM organization genes (GO:0030198) enriched in NT5E^+^ cells from all 9 cell lines. Cells ranked by ECM Organization gene set expression. E) RT-qPCR of select *NT5E-* associated markers in CD73-sorted cells (n=3, error bars=SEM, *p*-values = unpaired t-tests). F) Immunofluorescence staining of F-actin and Tenascin-C in CD73-sorted cells in 2D culture, 24 hours after FACS. Representative images of n=2.

**Supplemental Figure S3.**
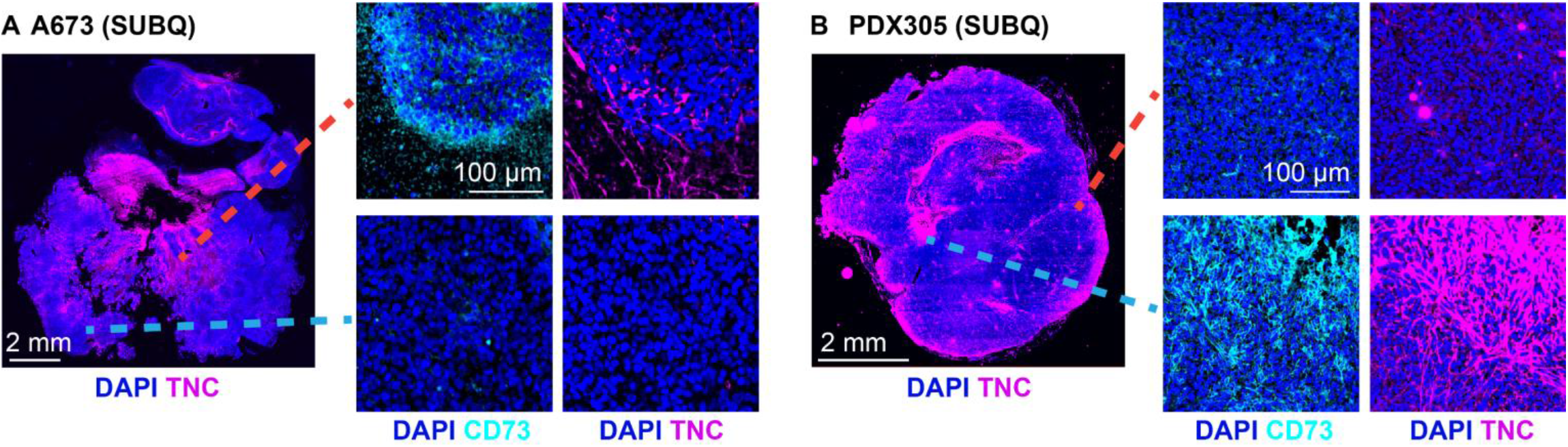
Spatial heterogeneity of CD73^+^ tumor cells and ECM deposition in additional xenograft and PDX-derived models. Related to Figure 3. A) Immunofluorescence for TNC and DAPI (nuclei) in an A673 subcutaneous (SUBQ) xenograft tumor section. Right, zoomed insets of CD73 and TNC immunofluorescence. B) Immunofluorescence for TNC and DAPI (nuclei) in a patient-derived xenograft (PDX305) subcutaneous tumor section. Right, zoomed insets of CD73 and TNC immunofluorescence.

**Supplemental Figure S4.**
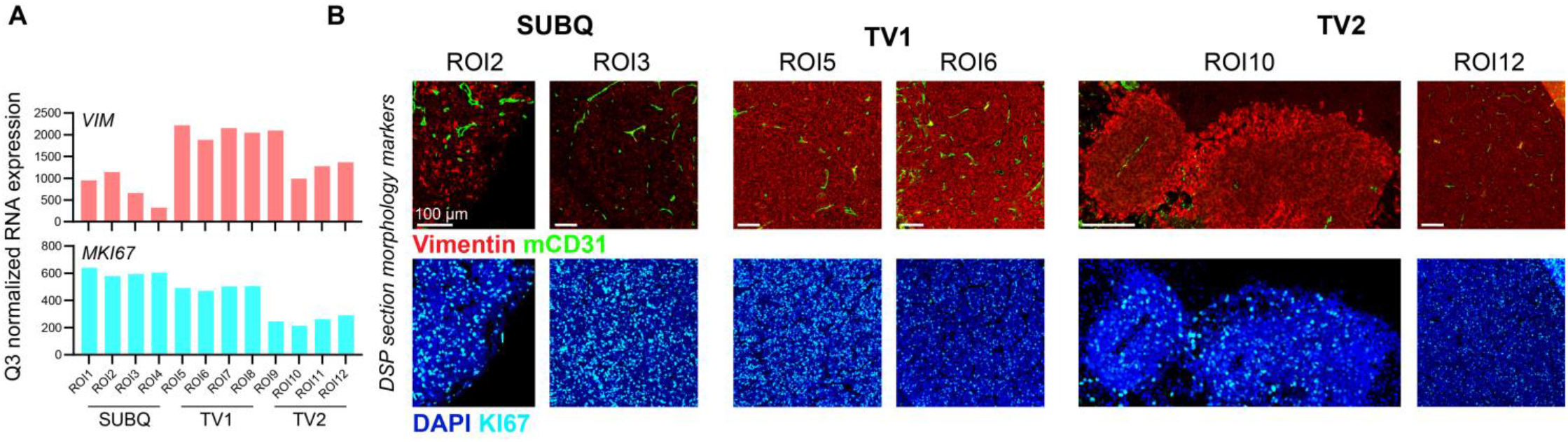
RNA vs. protein concordance in DSP ROIs. Related to Figure 4. A) Bar graphs depicting normalized probe expression of *VIM* and *MKI67* within each ROI subjected to DSP. B) Immunofluorescence of Vimentin, Ki67, mCD31, and DAPI on select DSP ROIs from each CHLA10 tumor.

**Supplemental Figure S5.**
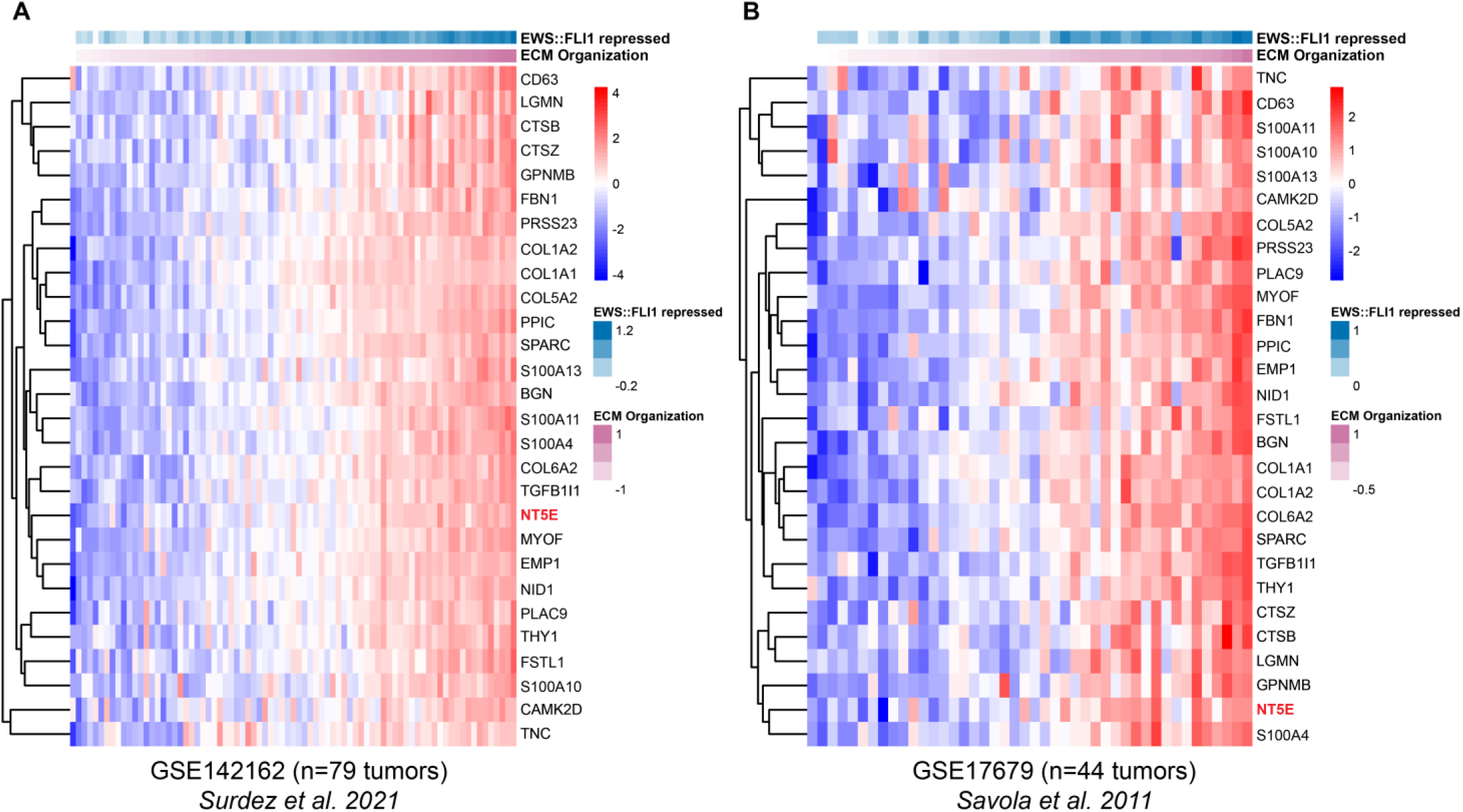
Patient tumor gene expression microarrays display heterogeneity of expression of *NT5E* and associated mesenchymal gene signature. Related to Figure 5. A-B) Heatmaps of *NT5E* and 27 associated marker gene expression vs. EWS::FLI1 repressed genes (27), ranked by ECM organization expression (A) GSE142162 (18) and (B) GSE17679 (37).

**Supplemental Figure S6.**
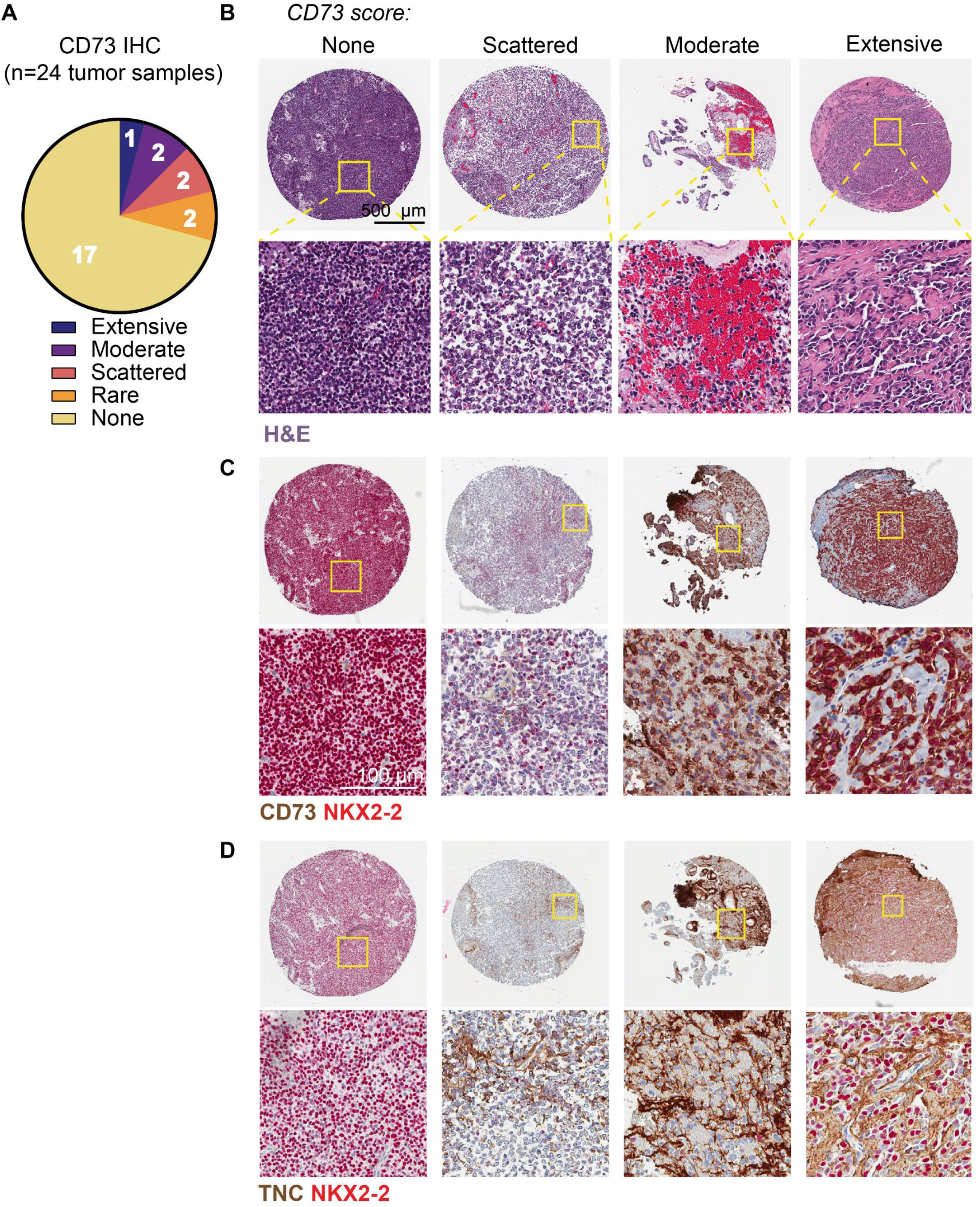
Intertumoral heterogeneity of CD73^+^ tumor cells and ECM deposition in patient biopsies. Related to Figure 6. A) CD73^+^ cell score for 24 EwS tumor biopsy specimens as determined by IHC score. Thirty-nine sections were scored from 24 tumor samples, representing 20 unique patients (details in Supplemental Table 5). B) Representative H&E images from negative, low, moderate, and high CD73 expressing tumors. C) & D) CD73 (C, brown) and TNC (D, brown) plus nuclear tumor marker NKX2-2 (red, C&D) IHC from TMA sections adjacent to H&E stained sections in (B).

## SUPPLEMENTAL TABLES

**Supplemental Table 1.** RNAseq-derived gene expression data for CD73^-^ and CD73^+^ A673 and CHLA10 cells.

**Supplemental Table 2.** Markers of *NT5E*^+^ cells from nine EwS cell lines by single cell sequencing (CITE-seq).

**Supplemental Table 3.** Q3 normalized gene expression counts and ROI information for CHLA10 xenograft digital spatial profiling.

**Supplemental Table 4.** List of genes that positively correlate with *NT5E* expression in 3 independent patient tumor cohorts and gene ontology associations.

**Supplemental Table 5.** Clinical information and CD73 IHC score for patient tumor tissue microarray.

**Supplemental Table 6.** Table of selected reagents and resources used.

## METHODS

### Cell lines

The EwS cell lines (A673, SKNMC, CHLA10, CHLA9, A4573, TC32, TC71, RDES) as well as U2OS, SW1353, and 293FT were obtained from ATCC and COG (https://www.childrensoncologygroup.org/) cell line repositories. A673, TC32, TC71, A4573, and SKNMC cell lines were maintained in RPMI 1640 media (Gibco) supplemented with 10% FBS (Atlas Biologicals) and 2mmol/L-glutamine (Life Technologies). RDES was cultured in in RPMI 1640 media supplemented with 15% FBS and 2mmol/L-glutamine. CHLA10 and CHLA9 were maintained in IMDM media (Fisher) supplemented with 20% FBS, 2mmol/L-glutamine, and 1X Insulin-Transferrin-Selenium-Ethanolamine (Gibco). U2OS was maintained in McCoy’s 5A media (Gibco) supplemented with 10% FBS and 2mmol/L-glutamine. H7-MSCs (kind gift from Dr. Sweet-Cordero) were maintained in alpha-MEM supplemented with 10% FBS, 2 mM L-glutamine, 1% antibiotic-antimycotic (Gibco). Cells were cultured at 37°C with 5% CO_2_. Cells were all confirmed to be mycoplasma free and identities subject to STR-confirmation every 6 months.

### PDX305 generation and characterization

PDX305 was generated from an abdominal soft tissue metastasis from 17-year-old male Ewing sarcoma patient. Cells were passaged as a patient-derived xenograft in NSG mice before cell line generation. PDX305 early passage cells were maintained in RPMI 1640 media (Gibco) supplemented with 10% FBS (Atlas Biologicals), 2mM L-glutamine (Life Technologies), and 1% Antibiotic-Antimycotic. Poly(A)-capture RNA-seq was performed on PDX305 cells as described in the RNA-seq sections. Data will be deposited to NCBI GEO upon publication.

### Flow cytometry and FACS

EwS cells were trypsinized, washed with PBS, resuspended in 2% FBS in PBS, and stained with fluorophore-conjugated antibodies (Supplemental table 6) for 30 min. on ice in the dark. Fluorophore-conjugated isotype controls and unstained controls were used for each experiment. Flow cytometry was performed on a BD Accuri C6 (10,000 events), with single cell identification by FSC and SSC. For FACS, cells were sorted on a BD FACSAria Il (see Supplemental Figure S1C for gating strategy). Analysis was conducted in FCS Express (De Novo Software).

### RT-qPCR

Total RNA was extracted from cells using the RNeasy Mini Kit (Qiagen) and cDNA was generated using the iScript cDNA Synthesis Kit (Bio-Rad) according to the manufacturer’s instructions. qPCR was performed using either Taqman Fast Universal PCR Master Mix (Applied Biosystems) assays or iTaq Universal SYBR-Green Supermix (Bio-Rad) on a Roche Light-Cycler 480 instrument (Roche Applied Science). Samples were run in technical triplicates and average Ct values were normalized to the geometric mean of two reference genes. The relative mRNA expression was calculated by the ddCt method. Primers and TaqMan assays listed in Supplementary Table S6.

### Tumor microarray preparation and immunohistochemistry

A tumor tissue microarray (TMA) of Ewing sarcoma tumors was prepared by Seattle Children’s Hospital Pathology (see Supplemental Table 5 for clinical details). For IHC, TMA sections were pretreated with CC1 Tris buffer (pH 8) for 32 minutes, primary antibodies (see Supplemental Table 6) were incubated 1:100 for 32 minutes at 36°C. Staining was detected using OptiView DAB (Roche).

### Immunofluorescence

#### Cells

For immunofluorescence, cells were fixed with 4% PFA in PBS for 10 min then washed with PBS. Sorted CD73^-^/CD73^+^ cells were plated at 60,000 cells/well in 8-well glass bottomed chambers overnight. Cells were permeabilized with 10 min. 0.3% Triton-X then blocked with 0.2% BSA in PBS for 1hr at room temperature. Primary antibodies were incubated for 1 hr at room temperature in 0.2% BSA (Supplemental Table 6). Alexa-fluor conjugated secondary antibodies were incubated for 1hr at room temperature in 0.2% BSA with 5% host serum (unless goat and donkey-derived secondary antibodies were used simultaneously, in which case host serum was not included). Signal was compared to matched species-specific IgG controls incubated at the same concentrations. For F-actin staining ActinGreen (ThermoFisher Scientific #R37110) was used at 2 drops/mL in the secondary antibody solution. DAPI was included during secondary antibody incubations to mark nuclei at 1 ug/mL. Coverslips were mounted with ProLong Gold and allowed to cure overnight at room temperature in the dark. Slides were imaged on a Leica SPE confocal at 40X magnification or scanned using an Olympus slide scanner (BX61) at 10X and 20X magnification.

#### Tissue

FFPE tumor samples from xenografts or primary patient samples were deparaffinized, then antigen retrieval was carried out using Diva Decloaker in a Biocare Medical decloaking chamber (120°C for 30 sec.) and slides were allowed to cool to room temperature. Slides were blocked with 0.2% BSA for 1 hour at room temperature. Primary and secondary antibodies (Supplemental Table 6) were incubated in 0.2% BSA for 1 hour at room temperature. When both goat and donkey derived antibodies were used, donkey-anti-goat secondary was applied first, washed 3x10 min. with PBS, then additional secondaries were added to prevent cross-species reactivity. DAPI was included during secondary antibody incubations to mark nuclei at 1 ug/mL. Coverslips were mounted with Prolong Gold and allowed to cure overnight at room temperature in the dark, then slides were stored at 4°C until imaging. Slides were imaged on a Leica SPE confocal at 40X magnification or scanned on an Olympus slide scanner at 10X and 20X magnification.

### Migration and Proliferation assays

Real-time cell analysis (RTCA) of cell migration and proliferation was monitored using a CIM-plate 16 or E-plate 16, respectively, on the xCELLigence DP system (Acea Bioscience). For proliferation, wells were coated with 0.2% gelatin and 100 μL complete media was added (5 x 10^3^ cells/well), and plates equilibrated for 1 hour at 37°C. For migration, before cell seeding, electrodes were coated with 0.2% gelatin and 50 μL complete media was placed in the upper chambers and 160 μL serum-free media was added to the lower chambers. 5 x 10^4^ cells/well were plated in the upper chamber in 100 μL media, then plates were equilibrated for 30 minutes at room temperature. Migration was evaluated up to 30 hours. Proliferation was evaluated up to 96 hours.

### 3D Collagen invasion assays

Spheroids were formed by plating EwS single cells at a density of 50,000 viable cells/mL in complete media in non-adherent 6 well plates overnight. Debris and single cells were removed by centrifuging samples for 10 sec. at 300g. Rat tail collagen (Gibco) was prepared at 1.8-2mg/mL in 0.08N acetic acid, supplemented with 10X DMEM (final concentration 1X), neutralized with 1M NaOH and allowed to polymerize on ice for 45-60 minutes until fibers formed (63). Spheroids and collagen gels were mixed and plated as 80 μL droplets and allowed to polymerize for 30 minutes at 37°C, at which point 1mL of complete IMDM media (20% FBS, 1X L-glutamine, ITS-X) was added. 40 μL underlays of polymerized rat tail collagen were first applied to the bottom of TC treated 24wp plates to prevent spheroids from attaching to or spreading along the plate surface. Spheroids were cultured 4-6 days then assessed by phase contrast microscopy and fixed for 15 minutes with 4% PFA in PBS. Invasion was scored as protrusive cells or multicellular strands extending from the spheroid border.

### Subcutaneous injections of CD73+ vs. CD73-cells

Cells were sorted based on CD73 positivity (see Supplemental Figure S1C) and 540,000 (CHLA10) or 720,000 (A673) CD73^-^ or CD73^+^ single cells were resuspended in PBS, diluted 1:1 in Matrigel (Corning), and injected subcutaneously into NSG mice (005557, Jackson Laboratory). Tumor formation and growth was measured by calipers every other day. All mice were euthanized at end point when the tumor of one animal reach 2cm diameter in any direction.

### Digital Spatial Profiling

GeoMx Digital Spatial Profiling (NanoString) was carried out according to manufacturer instructions. Tumor specimens were generated as follows: subcutaneous injection of 4x10^5^ CHLA10 cells in 50% Matrigel, primary tumors collected 54 days post-injection; tail vein injection of 7x10^5^ CHLA10 cells injected in PBS, tumors collected 28 days post-injection. Tumors were fixed for 24 hours in 10% non-buffered formalin before paraffin embedding. 3 sections of FFPE tumor tissues (1 from subcutaneous tumor, 2 from 2 different tail vein-injected mice) were sectioned and adhered to a single slide. Hybridization of the Human NGS Whole Transcriptome Atlas probe library was performed according to manufacturer instructions overnight at 37°C. Each probe is tagged with a photocleavable unique molecular identifier (UMI) for NGS-based readout. Slides were blocked for 30’ before staining with morphology markers (Supplemental Table 6), and SYTO DNA dye. After slide scanning, 12 ROIs comprising >1000 SYTO+ nuclei each were selected, UV-cleaved, and collected for NGS sequencing to quantify probe counts.

Library preparation and Illumina NGS were carried out by NanoString. The NanoString GeoMx NGS data analysis pipeline was used (version 2.3.4) to assess probe counts, including probe and count-level QC steps using the GeoMxTools R package. Gene level count data were normalized using the quartile 3 (Q3) method comparing the upper quartile of counts in each segment with the geometric mean of negative control probes. Highly variable genes were identified by ranking genes by coefficient of variation across all 12 ROIs (CV = standard deviation/mean).

### Western blots

Cells were lysed with RIPA buffer (Fisher Scientific) supplemented with protease and phosphatase inhibitors (Sigma). Western blot was performed using the Bio-Rad Mini-PROTEAN Tetra System. Following transfer, nitrocellulose membranes were blocked in Odyssey Blocking Buffer (LI-COR) for 1 hour. Membranes were washed and incubated rotating overnight at 4°C with a primary antibody (Supplementary Table #6). Membranes were then washed three times in TBST for 5 min each and incubated with a secondary antibodies (LI-COR IRDye 700CW or 800CW; 1:10,000) for 1 hour. Membranes were imaged on a LI-COR Odyssey scanner.

### Proteogenomic single-cell sequencing (CITE-seq)

CITE-seq performed as described previously (29). 500,000 cells were resuspended in PBS + 1% FBS and spun at 300g for 10 minutes at 4°C and then resuspended in 25 μL staining buffer (BioLegend #420201). 2.5 μL human TruStain FcX (BioLegend #42230) was added per sample and incubated at 4°C for 10 minutes. Hash-Tag Antibodies, used to label the samples and minimize batch effects, were added at 250 ng/sample (0.5 μL/sample) and incubated at 4°C for 30 minutes. Cells were washed twice in 1 mL staining buffer and spun at 300g for 5 minutes at 4°C. Samples were pooled 1:1 at 1000 cells/μL and libraries were generated using the 3’ V3 10X Genomics Chromium Controller following the manufacturer’s protocol (CG000183). Final library quality was assessed using the Tapestation 4200 (Agilent) and libraries were quantified by Kapa qPCR (Roche). Pooled libraries were then subjected to paired-end sequencing according to the manufacturer’s protocol (Illumina NovaSeq 6000). Bcl2fastq2 Conversion Software (Illumina) was used to generate de-multiplexed Fastq files and the CellRanger (3.1) Pipeline (10X Genomics) was used to align reads and generate count matrices. Further quality control and analyses were performed in R (see Lawlor Lab Github for code).

### Pathway analysis

Gene ontology and GSEA for selected gene lists was performed using MSigDB, Enrichr (64), Metascape (65), escape and dittoSeq (Bioconductor, see supplementary R notebooks on GitHub).

### RNA-sequencing

Poly(A)-capture RNA-seq was performed on PDX305 and FACS sorted CD73^-/+^ A673 and CHLA10 cells (RNA isolated from cell pellets after sorting). Libraries were prepared with NEBNext Ultra II RNA Library Prep kit and Paired end 150 bp sequencing was performed on a Novaseq600 by Novogene. Adapter trimming was performed using Trim Galore (Babraham Institute). Trimmer reads were aligned to GRCh38 using the STAR aligner (29). Counts and differential expression analysis were calculated by DESeq2 v1.18.1 using FDR-adjusted *p*-values <0.05.

### Public data sources

Log_2_ expression values for tumor microarray datasets (GSE34620, GSE142162, GSE17679) (18, 36, 37) were downloaded from the R2 Genomics Analysis and Visualization Platform. Only Ewing sarcoma patient tumor samples were included.

### Statistics and software

Statistical tests were conducted in GraphPad Prism 9. Error bars = SEM unless otherwise specified. Unpaired t-tests were used unless otherwise specified. ns=p>0.05, *=p<0.05, **=p<0.01, ***=p<0.001, ****=p<0.0001. BioRender.com was used to prepare experimental schematics.

### Study approval

All experiments in conducted in mice were approved by the Institutional Animal Care and Use Committee of Seattle Children’s Research Institute.

### Data and Code Availability

Sequencing data generated in this study will be deposited to NCBI GEO upon publication. Analyses in R will be available upon publication at https://github.com/LawlorLab/.

## Author contributions

AA & EW designed and conducted experiments, analyzed and interpreted data, drafted and edited the manuscript. AA, EW, NG, VH, SB, SK, and OW designed, conducted, and analyzed experiments. ER and XD generated and performed IHC on the tumor microarray samples. ST, OW, and SF performed bioinformatics analyses. SS, RV, WJ and EN generated and characterized PDX305. AA, EW, and EL wrote and edited the final manuscript, with input from all authors. EL supervised the study and secured funding. Co-first authors were ordered randomly.

## Acknowledgments

The authors thank members of the Lawlor lab for helpful discussion, and the staff at the Fred Hutchinson and Seattle Children’s Research Institute flow cytometry, comparative medicine, and bioinformatics cores. Grant and gift support for this work is gratefully acknowledged and was provided by the following sources: NIH/NCI R01 CA215981 (ERL), F31CA247104 (AA); AACR-QuadW Sarcoma Fellowship in Memory of Willie Tichenor (EDW); Sam Day Foundation (ERL); the 1M4Anna Foundation (ERL, SNF); Lafontaine U-Can-Cer-Vive Foundation (ERL). This research was supported by the Flow Cytometry Shared Resource of the Fred Hutch/University of Washington/Seattle Children’s Cancer Consortium (P30 CA015704).

